# A strong priority effect in the assembly of a specialized insect-microbe symbiosis

**DOI:** 10.1101/2024.04.26.591361

**Authors:** Jason Z. Chen, Anthony Junker, Iris Zheng, Nicole M. Gerardo, Nic M. Vega

## Abstract

Microbial community assembly is determined in part by interactions between taxa that colonize ecological niches available within habitat patches. The outcomes of these interactions, and by extension the trajectory of community assembly, can display priority effects - dependency on the order in which taxa first occupy these niches. The underlying mechanisms of these phenomena vary from system to system and are often not well resolved.

Here, we characterize priority effects in colonization of the squash bug (*Anasa tristis*) by bacterial symbionts from the genus *Caballeronia*, using pairs of strains that are known to strongly compete during host colonization, as well as strains that are isogenic and thus functionally identical. By introducing symbiont strains into individual bugs in a sequential manner, we show that within-host populations established by the first colonist are extremely resistant to invasion, regardless of strain identity and competitive interactions. By knocking down the population of an initial colonist with antibiotics, we further show that colonization success by the second symbiont is still diminished even when space in the symbiotic organ is available and physically accessible for colonization. We propose a paradigm in which resident symbionts exclude subsequent infections by manipulating the host environment, partially but not exclusively by eliciting tissue remodeling of the symbiont organ.

**Importance:** Host-associated microbial communities underpin critical ecosystem processes and human health, and their ability to do so is determined in turn by the various processes that shape their composition. While natural selection acts on competing genotypes and species during community assembly, the manner by which selection determines the trajectory of community assembly can differ depending on the sequence by which taxa establish within that community. We document this phenomenon, known as a priority effect, during experimental colonization of a North American insect pest, the squash bug *Anasa tristis*, by its betaproteobacterial symbionts in the genus *Caballeronia*. Our study demonstrates how stark, strain-level variation can emerge in specialized host-microbe symbioses simply through differences in the order by which strains colonize the host. Understanding the mechanistic drivers of community structure in host-associated microbiomes can highlight both pitfalls and opportunities for the engineering of these communities and their constituent taxa for societal benefit.

## Introduction

The composition of microbial communities is determined not just by colonization and extinction events among its constituent members, but also by the timing of these events. Cases where colonization ability of later-arriving taxa are conditional on the taxa already established in that habitat are referred to as showing *priority effects*. When priority effects manifest during microbial community assembly, such communities can reach alternative states contingent on the history and timing of past colonization events.

Microbial community assembly across different systems may exhibit priority effects for many reasons(1, 2). Characterizing the mechanisms underlying priority effects (3–7) is of interest not only because priority effects are widespread and striking ecological phenomena (1), but also because of the potential to engineer communities via control of the order of introduction. The most simple mechanism underlying a priority effect may be when early colonizers occupy a physical or nutritional niche, leaving fewer resources for later colonizers. In addition, early colonizers can modify the physical or chemical environment in a way that facilitates or hinders colonization by immigrating taxa (3, 8). Finally, already-resident populations can deploy positively density-dependent strategies, such as socially-mediated transcriptional regulation (9) and collective defense in spatially structured habitats (10–14), to prevent invasion by less numerically dominant competitors.

Microbial communities are frequently associated with multicellular hosts (15). Host-associated microbial communities follow the same rules of assembly as any other ecological community (16). As living habitats, however, hosts also impose dynamic, top-down effects on community assembly. When symbiotic microbes colonize host organs, they stimulate immunological, physiological, developmental, and behavioral responses by the host (17, 18), affecting the probability of establishment by later-arriving microbes. Thus, symbionts can rapidly modify an ecological niche by eliciting transitions in the state of the host, which can impose a top-down priority effect.

The squash bug *Anasa tristis* hosts extracellular betaproteobacterial symbionts of the genus *Caballeronia* (Dobritsa et al., 2017; Dobritsa & Samadpour, 2016; Peeters et al., 2016), which it relies on for proper growth and development (22). Bacteria are housed in a specialized region of the midgut, the M4, which acts as the symbiotic organ and in which *Caballeronia* spp. are the dominant taxa. The M4 in *A. tristis* consists of two rows of numerous crypts whose primary function appears to be the maintenance of symbiont biomass. Presence and morphology of the symbiont organ are strongly conserved among related insects (the Pentatomomorpha) (23), which maintain beta- and gamma-proteobacterial symbionts such as *Caballeronia* (24–31). Although this symbiosis is highly specific, many genetically distinct *Caballeronia* symbiont isolates are capable of colonization in *A. tristis*, with no apparent difference in the fitness benefits that they confer to the host (33, 34). There is also striking heterogeneity between individual squash bugs in the strain-level taxonomic composition of their symbiont communities (34), raising the question of how community structure and strain diversity emerges and is maintained (35).

The bean bug *Riptortus pedestris* of the family Alydidae, which last shared a common ancestor with *A. tristis* over 100 million years ago (23, 36, 37), has yielded remarkable insights into the establishment and maintenance of specialized *Caballeronia* symbioses in insects (32). Symbiotic infection occurs through a physical bottleneck in the midgut at the anterior entrance to the M4, called the constricted region. In *Riptortus*, this constricted region imposes selectivity on the bacterial cells that can access the symbiotic organ (38). Once symbionts have passed through the constricted region and colonized the M4, the constricted region of *Riptortus* closes off entirely, preventing food intake into the M4 for the duration of host development and rendering the midgut physically discontinuous in nymphs (38, 39). The closure of the constricted region within 12-15 hours after establishment of early-arriving symbionts has been implicated in preventing subsequent symbionts from colonizing the M4 (40). Thus, at least in *Riptortus*, gut anatomy is remodeled in a way that imposes a priority effect on community assembly, preventing compatible symbiont strains from invading established symbiont populations within the host. While the presence of the constricted region is conserved among host insects in this clade (38), the degree to which it plays similar roles in *Caballeronia*-associated insects other than *Riptortus* is unknown.

Here, we evaluate the evidence for priority effects, and the mechanisms underlying them, in symbiont colonization of the *A. tristis* symbiotic organ. Using two strains that exhibit an asymmetrically competitive interaction, we show that a priority effect exists that allows the first symbiont strain to completely exclude establishment of a second strain regardless of competitive ranking. This phenomenon emerges even when two strains are isogenic, suggesting that the priority effect takes place regardless of strain identity or the nature of between-strain interactions. By depleting symbiont populations with antibiotics and by examining the timing of gut development after colonization, we rule out the role of spatial occupancy and symbiont-elicited tissue remodeling as mechanisms governing priority effects in the squash bug-*Caballeronia* system.

## Results

### Priority effects override between-strain competition

To assess whether priority effects play an important role in the establishment of symbiont populations in *A. tristis*, we sequentially inoculated cohorts of nymphs with two unrelated bacterial strains, *C. zhejiangensis* GA-OX1 and *C. sp. nr. concitans* SQ4a (Figure 1A), each expressing a green or red fluorescent protein marker (GFP or RFP, respectively) that represent different clades within the bacterial genus *Caballeronia*. In previous experiments in which bugs were co-inoculated with large doses of both strains, GA-OX1 tended strongly to outcompete SQ4a (35). If priority effects play a role in community assembly in this symbiosis, the early-arriving colonist (referred to here as the resident strain) should dominate symbiont communities regardless of which strain or fluorophore is used. Conversely, if community assembly is driven primarily by competition between colonizing strains, then the order of colonization should not be significant, and GA-OX1 should tend to exclude SQ4a, as observed when both strains colonize the host simultaneously (35). When bugs were inoculated first with GA-OX1 GFP, and then with SQ4a RFP, all bugs were colonized exclusively with the first colonist, GA-OX1 GFP (Figure 1B, top); SQ4a RFP was not recovered from any of these nymphs (Figure S1A, left). Conversely, when bugs were inoculated first with SQ4a RFP and then with GA-OX1 GFP, all bugs were colonized exclusively with the first colonist, SQ4a RFP (Figure 1B, bottom; Figure S1A, right), despite its documented competitive disadvantage during simultaneous pairwise competition with GA-OX1 (35). Thus, symbiont composition between these two treatments significantly differed according to colonization order but not according to strain identity (Figure S1A, W = 0, p-value = 9.98×10^-13^). We observed this effect regardless of which fluorescent marker was associated with which strain (Figure 1B, 1C). Our results demonstrate a strong priority effect producing mutual superinfection exclusion between *Caballeronia* symbiont strains during insect colonization.

**Figure 1.**
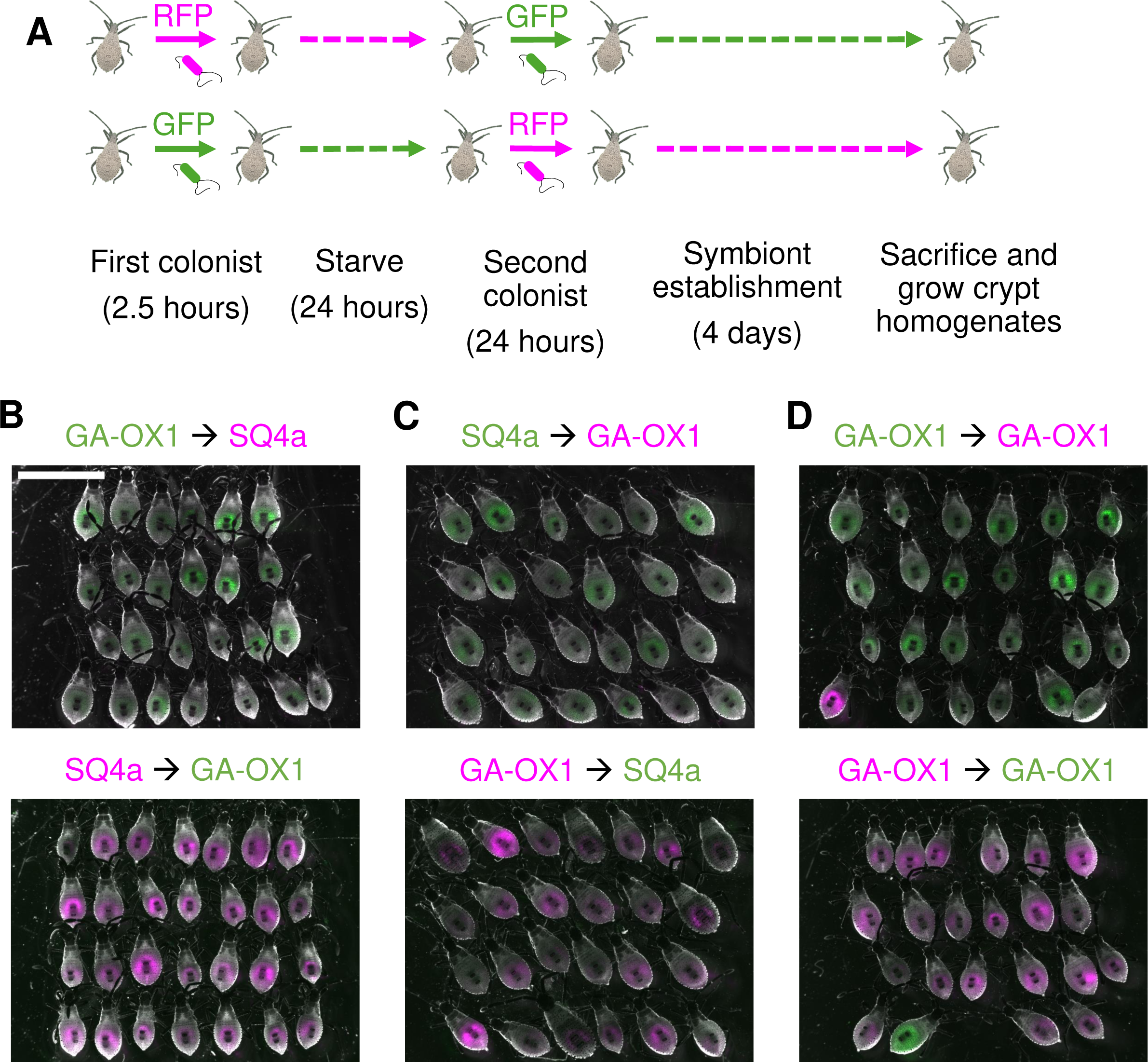
Random colonization order mediates the outcome of colonization. Experimental design. Nymphs were starved overnight, then allowed to feed on a 1:100 dilution of either a GFP-expressing or RFP-expressing symbiont strain (first colonist) for two hours before being starved for 24 hours. The next day, nymphs were fed with the strain of the opposite color *ad libitum* for 24 hours (second colonist). Infections were allowed to establish for four days before nymphs were sacrificed. (B-D) Sequential colonization by counter-labeled symbiont strain pairs. Strain identity and fluorophore of the first and second colonists are shown above each plot. B. Sequential colonization by GA-OX1 GFP and SQ4a RFP. C. Sequential colonization by GA-OX1 RFP and SQ4a GFP. D. Sequential colonization by isogenic GA-OX1 GFP and GA-OX1 RFP.

### Priority effects exclude superinfection by isogenic strains

To further explore the possible effects of inter-strain competition, we repeated these experiments using a neutral competition scenario where hosts were colonized sequentially with counter-labeled, isogenic isolates of GA-OX1. If asymmetric competitive interactions between colonizing symbionts are necessary for priority effects, then colonization order should not be important in this case, and individual hosts should be co-colonized with GFP+RFP isogenic symbionts. Conversely, if priority effects operate independently of competitive interactions between symbiont strains, then priority effects should manifest as in the previous experiment. As before, outcomes differed based on colonization order (Figure S1B, W = 23.5, p-value = 1.53×10^-9^). Individual hosts were colonized almost exclusively with the first colonist, regardless of whether GA-OX1 GFP (Figure 1D, top; Figure S1B, left) or RFP (Figure 1D, bottom; Figure S1B, right) was presented first. In each of these treatments, exactly one bug was exclusively colonized with the counter-labelled second strain rather than the first-inoculated strain. Because these two nymphs did not contain mixed infections, we hypothesize that the first strain failed to successfully establish in a small fraction of hosts, allowing the second strain to colonize as if no other strains had come before it; this is consistent with prior results indicating a non-zero failure rate for initial colonization even by beneficial microbes within their native hosts (41, 42). Our results demonstrate that the strong priority effect in symbiont colonization that we observed in the previous experiment cannot be explained as a result of direct conflict between genetically distinct microbial competitors.

### Priority effects are not due to pre-emption of space within the gut

Following the example of previous research in the alydid bean bug *Riptortus* (40), we next interrogated some possible mechanisms that might be responsible for the strong priority effect that we observed. One possible mechanism is competitive exclusion as a result of spatial occupancy of the symbiotic organ, in which full occupancy of the crypts by the first microbe could reduce the physical space available for colonization by the second strain. This has been demonstrated in other systems (42), including those in which microbes colonize analogous crypt-like spaces consisting of a small, confined lumen with only one opening (Conwill et al., 2022; Whitaker et al., 2017) but see (6)). This hypothesis implies that purging the symbiotic organ of previously established residents should restore efficient establishment by the second colonizer.

To knock down intra-host populations of a colonizing symbiont, we adapted a protocol (40) to clear hosts of symbionts using trimethoprim (Tp) (Figure 2A), an antibiotic to which GA-OX1 is sensitive *in vitro.* We tested the efficacy of Tp *in vivo* by treating nymphs pre-colonized with GA-OX1-RFP with the antibiotic, then dissecting the symbiotic organs to determine clearance of the symbiont. After Tp treatment was over, we moved the nymphs to their normal, antibiotic-free diet to purge as much Tp as possible and to permit recovery of any symbionts which still might be in the symbiotic organ (Figure 2A). As observed in *Riptortus* (40), we found that even after an extended recovery period of three days, many nymphs exhibited diminished fluorescence compared to colonized nymphs not treated with antibiotics (Figure 2B). However, closer examination found individual RFP puncta, suggestive of intact GA-OX1 RFP cells, inside of symbiotic organs dissected from all six individuals we chose for microscopy (Figure 2C). These results indicate that low-level occupancy of the symbiotic organ by potentially viable RFP symbiont cells persists in spite of antibiotic treatment, but that the bulk of the symbiont population is nonetheless cleared and/or its growth suppressed nonetheless.

**Figure 2.**
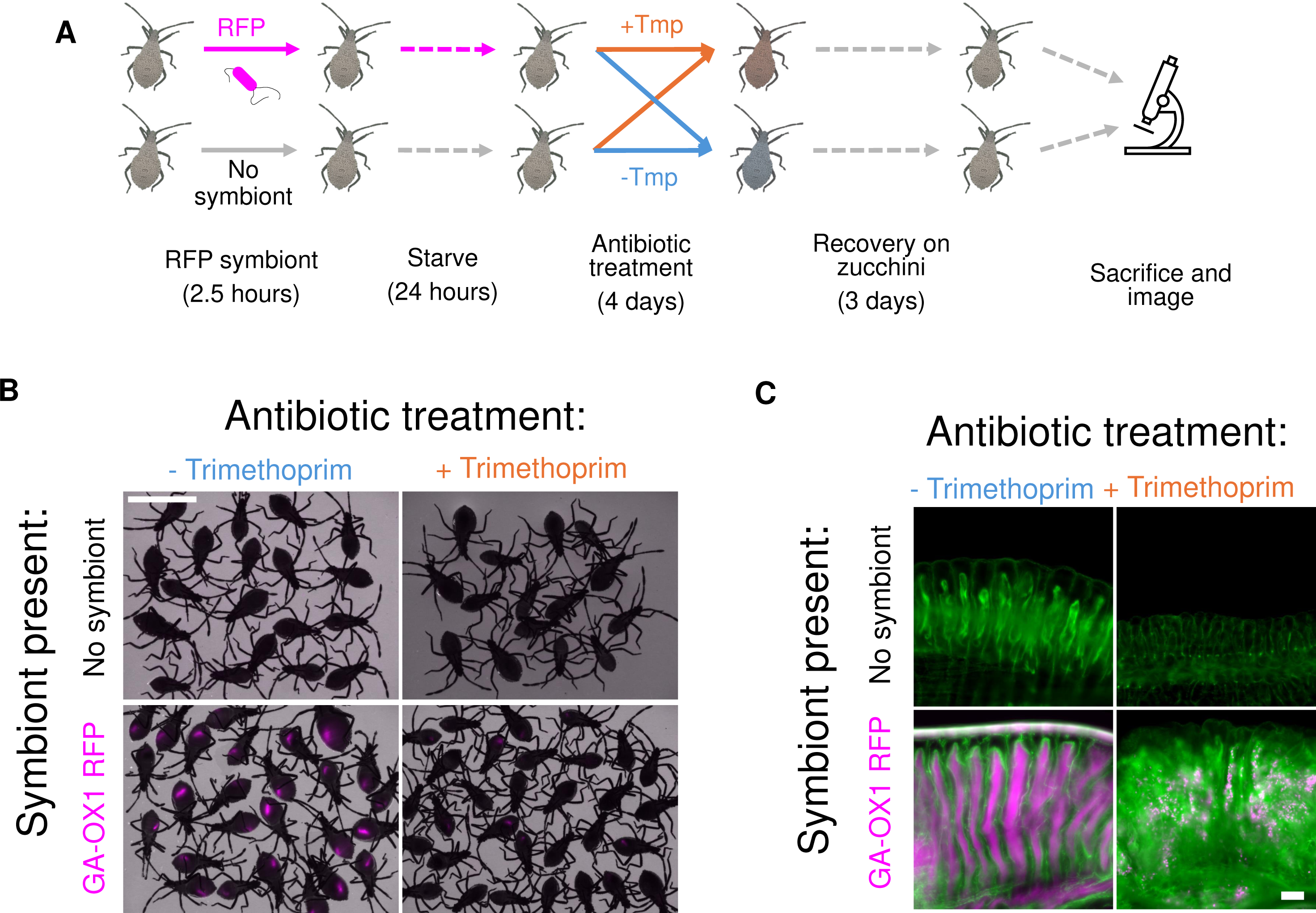
Trimethoprim suppresses symbiont proliferation in squash bug nymphs after colonization. A. Experimental design. Nymphs were fed for only 2.5 hours with either sterile inoculation medium or with inoculation medium containing either GA-OX1 RFP before being starved for 24 hours. The next day, aposymbiotic or inoculated nymphs were fed either sterile inoculation medium or inoculation medium containing >100 µg/mL trimethoprim (Tp). Bugs were maintained on this diet for four days before being moved to squash to allow potential recovery of any surviving symbionts. On the last day of recovery, whole nymphs were imaged to confirm successful depletion of RFP symbionts, and the individuals with the dimmest RFP signal were dissected for widefield microscopy. B. Colonization of an antibiotic sensitive strain after dietary administration of Tp. Top row: Symbiont-free nymphs never exposed to antibiotics (left) or treated with Tp for four days (right). Bottom row: Nymphs previously colonized with GA-OX1 RFP that were either never exposed to antibiotics (left) or treated with Tp for four days (right). Scale bar = 5 mm. C. Dietary Tp does not totally eliminate antibiotic-sensitive symbiont strains. Top: Crypts from symbiont-free nymphs never exposed to antibiotics (left) or treated with Tp for four days (right). Bottom: Crypts from nymphs previously colonized with GA-OX1 RFP that were either never exposed to antibiotics (left) or treated with Tp for four days (right).

Despite our inability to fully eradicate symbionts with Tp, symbiont density remained suppressed in the M4 for many days after Tp administration ceased, suggesting persistent post-treatment effects of the drug in vacating the symbiotic organ. We tested the hypothesis that this vacant space could be available for colonization. To ensure that any drug remaining after Tp treatment would not interfere with inoculation, we used an RFP-expressing, Tp-resistant (45) (dsRed-TpR) symbiont as the second colonist (Figure 3). We inoculated bugs with GA-OX1 GFP, then treated them with Tp to knock down the first colonist. After confirming the successful knockdown of GFP signal in live nymphs, we inoculated them with a high density of GA-OX1 dsRed-TpR for 24 hours (Figure 3A). As we expected, GA-OX1 dsRed-TpR consistently colonized symbiont-free nymphs regardless of whether they were previously treated with Tp (Figure 3B, top row); this provided a baseline for expected colonization by this strain. In line with our previous results, the second strain did not colonize nymphs in which GA-OX1 GFP was never suppressed with Tp (Figure 3B, bottom left). Surprisingly, GA-OX1 dsRed-TpR nearly always failed to colonize even after curing insects of the Tp-sensitive GFP strain; instead, almost all the nymphs either recovered GFP or remained non-fluorescent (Figure 3B, bottom right) a full week after their last antibiotic exposure. In these experiments, antibiotic treatment did not produce a significant increase in colonization by a second, resistant strain, despite providing it an ostensible fitness advantage over the susceptible resident strain (binomial regression: p_GFP_first_ = 6.58×10^-7^; p_Tp_cure_= 0.37). Our results indicate that the vacant space left in the M4 by antibiotic suppression of resident symbionts is not accessible for colonization by a second, incoming strain.

**Figure 3.**
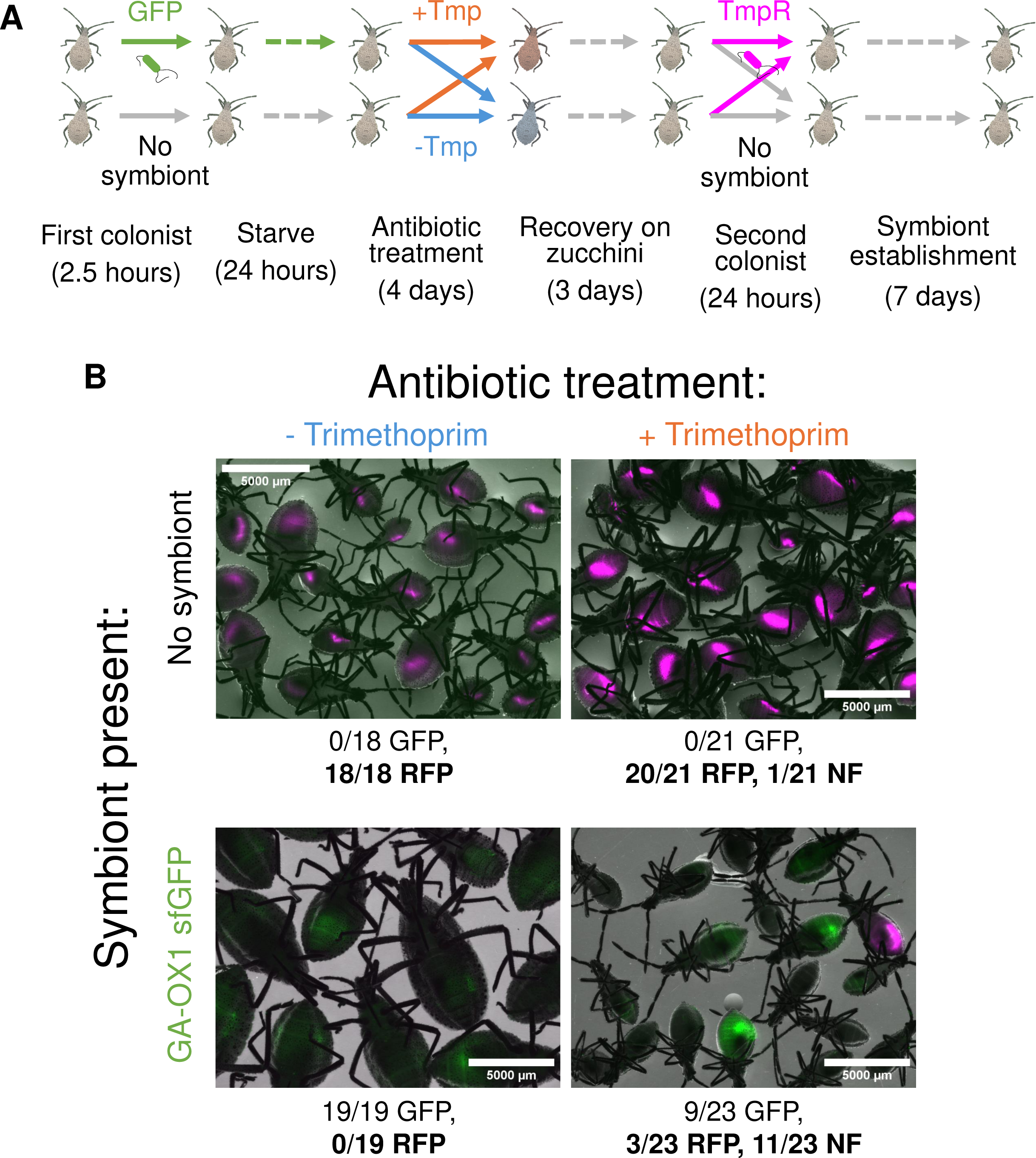
Priority effects after antibiotic depletion of the first colonizing symbiont strain. A) Experimental design. Nymphs were fed either sterile inoculation medium or inoculation medium containing GA-OX1 GFP for 2.5 hours, before being starved for 24 hours. The next day, aposymbiotic or inoculated nymphs were fed either sterile inoculation medium or inoculation medium containing >100 µg/mL Tp. Bugs were maintained on this diet for four days before being moved to squash to allow recovery of any surviving symbionts. On the last day of recovery, nymphs were imaged to confirm the successful depletion of GFP symbionts. Next, bugs in each treatment were fed either sterile inoculation medium or inoculation medium containing GA-OX1 dsRed-TpR for 24 hours. Nymphs were thereafter maintained on squash for one week to permit RFP symbiont colonization and further GFP symbiont recovery until sacrifice and imaging. B) Colonization by a Tp-resistant symbiont strain after clearance of a susceptible resident symbiont population. Top row: GA-OX1 dsRed-TpR colonization of aposymbiotic nymphs which had never been exposed to Tp (left) or had been treated with Tp for four days (right). Bottom row: GA-OX1 dsRed-TpR colonization of nymphs, inoculated previously with GA-OX1 GFP, which had never been exposed to antibiotics (left) or had been treated with Tp for four days (right). Bugs that exhibited neither RFP nor GFP fluorescence were scored as non-fluorescent (“NF”).

### Priority effects cannot be explained by symbiont-elicited tissue remodelling of the symbiotic organ

In *Riptortus* (40), ingestion of the first symbiont strain only prevents colonization of the second strain after a 15-hour interval between inoculations. This accompanies the physical closure of the entrance to the symbiotic organ, known as the constricted region, at around the same time. This closure seems to depend on the presence of a colonizing symbiont in the host (40). The constricted region is highly conserved in pentatomomorphan insects with beta- and gamma-proteobacterial gut symbionts (38). The constricted region is likewise present in squash bug nymphs (Figure 4A) and forms a closed plug without an apparent lumen after successful *Caballeronia* colonization (Figure 4B and 4C).

**Figure 4.**
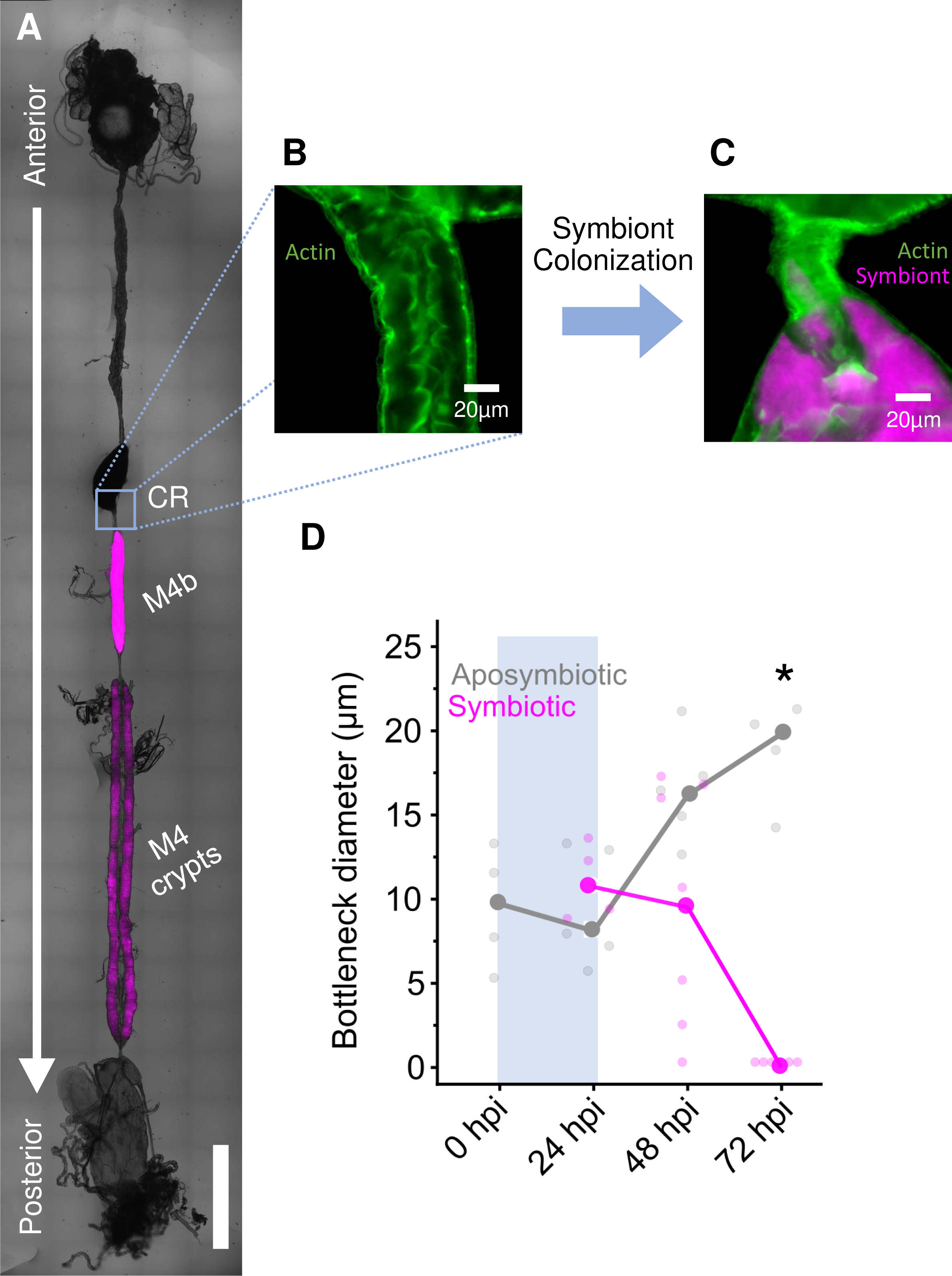
Closure of the constricted region (CR) within the gut is not necessary for the priority effect. A. Tilescan of the entire midgut of a fifth instar squash bug nymph, linearized to depict the sections of the midgut. The magenta signal localized to the M4 crypts and the M4b is indicative of colonization with GA-OX1 RFP. The CR is indicated in the orange box. B. (inset) The CR of an aposymbiotic second instar nymph, with an open bottleneck. Host actin is stained in green. C. Closed CR of a second instar nymph colonized with GA-OX1 RFP (magenta). D. Changes in the diameter of the bottleneck within the CR of second instar nymphs kept symbiont-free (gray) vs. colonized with GA-OX1 (magenta), over the course of 72 hours. Bugs colonized with symbionts exhibit a reduction in the size of the bottleneck between 24 and 72 hours post infection (hpi), while aposymbiotic bugs do not, consistent with a slow tissue-remodeling response to symbiont colonization. However, this reduction in the diameter of the lumen does not take place within the 24 hour period in which the priority effect emerges (shaded in light blue-gray) in our experiments. Smaller faded points indicate exact measurements, while large points and lines indicate averages for each treatment.

We reasoned that, as in *Riptortus*, the closure of the constricted region in the squash bug could explain the inability of the second strain to colonize the host, despite the availability of physical niche space in the symbiotic organ after antibiotic treatment. In other words, the priority effect we observed in all our experiments might result not from direct microbe-microbe interactions, but rather from microbial modulation of host physiology. We suspected that the continued presence of bacterial cells in symbiont-colonized nymphs fed Tp (Figure 2C, bottom right) could be sufficient to trigger the closure of the constricted region, just as in fully symbiont-colonized bugs. This could physically prevent antibiotic-resistant symbionts from entering the symbiotic organ despite the low density of antibiotic-susceptible cells resident within (Figure 3B, bottom right). Thus, we hypothesized that the constricted region must be completely closed in nearly all squash bug nymphs within 24 hours to explain the prevalence and magnitude of the priority effects in our experiments (Figure 1).

To ascertain whether the closure of the constricted region could impose a physical barrier to subsequently ingested symbionts, we measured the inner diameter of the lumen within the constricted region every 24 hours post-symbiont inoculation for three days. We compared the change in diameter of the lumen in symbiont-colonized nymphs with those of aposymbiotic nymphs over the same time period. Contrary to our expectations, we observed no significant differences in the luminal diameter of the constricted region 24 hours post infection, although the robust priority effect we observed in our previous experiments (Figure 1) is in place sometime before this point (Figure 4D, Wilcoxon signed-rank test; p_24hpi_= 0.39). Instead, the constricted region appears to fully close between 48 to 72 hours post infection (p_48hpi_=0.19; p_72hpi_= 5.7×10^-3^). We conclude that, at least in our model system, the timescale at which closure of the constricted region takes place is not sufficiently rapid to explain the complete exclusion of the second symbiont from colonization.

## Discussion

In previous work, we demonstrated that communities of *Caballeronia* bacterial symbionts in the squash bug *Anasa tristis* are highly variable between hosts (34) and that transmission bottlenecks during symbiont colonization are sufficient to drive these patterns (35). In this work, we demonstrate a strong, rapid priority effect in colonization of the squash bug *Anasa tristis* by symbiotic bacteria in the genus *Caballeronia*. By manipulating the order by which colonization takes place within a 24-hour time window, we show that the first strain to colonize a host prevents colonization by the second. This effect is agnostic to symbiont strain and cannot be reversed by making physical space available for colonization. We argue that symbiont communities in *A. tristis* are completely closed off from microbial immigration soon after their establishment, and community assembly thereafter is governed by internal processes, i.e. drift, selection, and mutation (16). Thus, rapid priority effects maintain heterogeneity in population-level microbial community composition, as described in other systems (e.g. (4)), by insulating communities from the homogenizing effects of between-host dispersal.

Priority effects manifest in populations and communities across scales and systems (46), but the underlying mechanisms are often unknown. We first showed that strain identity is not important; two different host-beneficial strains mutually prevented each other from subsequent colonization, and isogenic lineages of the same strain showed the same effect. Next, we demonstrated that the availability of space within the M4 does not mean that this space can be colonized by a subsequent symbiont, implying that the open space in the symbiont organ is somehow inaccessible. Finally, we determined that host tissue remodeling and closure of the M4, which is normally elicited by symbiont colonization, does not occur rapidly enough after colonization to exclude subsequent symbionts within the timeframe of our experiments.

Our results partly corroborate previous work in *Riptortus pedestris*, which also exhibits strong priority effects in *Caballeronia* colonization (40). However, while Kikuchi *et al.* (2020) attribute this priority effect to a concomitant narrowing and closure of a physical constriction in the midgut, we were unable to identify the mechanism(s) responsible for priority effects in *Anasa tristis*. Because of methodological differences between these studies, future work should examine the timing of these important developmental events across a range of bug-*Caballeronia* symbioses.

There are many potential mechanisms that we could not explore due to the limitations in the experimental tractability of our system. First, we were unable to completely eliminate resident symbionts in the symbiotic organ with antibiotic treatment, contrary to previous reports (40). However, the bacterial populations remaining after Tp treatment were visibly small and sparse, and we consider it unlikely that interactions of these remaining cells with subsequent colonists, including niche interactions and warfare (9, 47), would be sufficient to entirely prevent entry of a second colonist.

Other mechanisms on the part of the host might mediate the priority effect. First, although microscopy did not reveal a reduction in the physical diameter of the constricted region, the entrance to the M4, it is possible that changes in the secretions in this region might block off access to subsequent colonization. Such modifications would be invisible to the actin stain we used to visualize the lumen of the constricted region. Furthermore, we suspect that colonization likely causes shifts in immunity or the nutritional environment throughout the gut, which may pose a more rapidly erected barrier to superinfection than tissue remodeling in the constricted region. Future studies should address both microbial and host contributions to the priority effect that we observed in this study.

Our results have applied implications. Strong priority effects could be exploited to permanently colonize squash bugs with symbiont strains that express desirable phenotypes in the field. This is a long-term goal of many “paratransgenic” approaches in insect livestock and pests (48–51). Possible applications include use of engineered symbionts that reduce pest fitness(52), suppress pathogen vectoring (53), or express a kill switch that terminates symbiont infection of hosts. Engineered symbionts in the *Anasa-Caballeronia* system may be useful as part of a broader strategy for integrated pest management, even if they have decreased fitness relative to wild strains, as long as they are the first to colonize their hosts. Further studies to determine the feasibility of symbiont-mediated pest management using this strategy should further investigate the molecular mechanism(s) behind priority effects in this system to fully take advantage of this property in the assembly of these host-beneficial communities.

## Materials and Methods

### A) Study system

Adult insects were maintained at 28-30°C on yellow crookneck squash plants (*Cucurbita pepo* ‘Goldstar’) housed in 30 cm^3^ mesh cages. Eggs were routinely collected and surface-sterilized by rinsing sequentially in 70% ethanol, 20% bleach, and sterile water. Upon hatching, symbiont-free nymphs were maintained on surface-sterilized, parafilmed organic zucchini slices until they were ready to be used in inoculation experiments. Symbiont-free nymphs molt to the second instar two days after hatching from eggs, whereupon they become competent for symbiont colonization. Nymphs used in inoculation experiments were never more than one week old.

*Caballeronia zhejiangensis* GA-OX1 and *Caballeronia sp. nr. concitans* SQ4a are isolates from wild-caught squash bugs in northeast Georgia (22, 34, 54). These strains represent two different clades (*sensu* (21)) within the genus *Caballeronia* but exhibit no distinguishable differences in the degree of host benefit conferred (22, 34). For experiments, both GA-OX1 and SQ4a were cultivated in the laboratory at 25°C on nutrient agar (NA; 3 g/L yeast extract, 5 g/L peptone, 15 g/L agar) plates or in low-salt Luria-Bertani Lennox broth (LB; Sigma-Aldrich L3022).

### B) Sequential inoculation experiments

The generalized experimental design is depicted in Figure 1A. *Caballeronia* strains were previously labelled with superfolder GFP (“GFP”) or dTomato (“RFP”) (35, 55) using the mini-Tn7 system (56, 57). Fluorescent proteins are genomically integrated at a specific, neutral intergenic site and are thus stably maintained with minimal effects on phenotype and fitness *in vitro* (58–60) and with minimal opportunity for horizontal gene transfer.

Fluorescently labelled *Caballeronia* strains were streaked out on NA plates and grown for two days at 25°C. Liquid cultures were started by picking single colonies into 2 mL LB and incubated at 25°C overnight in glass tubes shaking at 200 rpm. Three hundred µL of each culture was diluted into 1000 µL 1X phosphate buffered saline (PBS) in a 1.5 mL microcentrifuge tube and washed by pelleting at 10,000 x g for two minutes at 4°C, removing the supernatant, and resuspending in 1000 µL PBS. After a second wash step, the pellet was resuspended in 300 µL PBS.

In preparation for the first round of inoculation, nymphs were starved for 20-24 hours in a surface-sterilized plastic rearing box, supplied with only 100 µL sterile water spotted as droplets across the lid of the box with a 1000 µL tip for hydration. The next day, the starved nymphs were transferred into another clean, surface-sterilized plastic rearing box. For each *Caballeronia* strain, an inoculum was prepared by diluting 2 µL of washed live cells into 200 µL of a defined inoculation medium, an aqueous solution containing 2% w/v glucose and 10% v/v PBS. Droplets totaling 150 µL of inoculum were spotted onto the plastic surface of the box, and nymphs were allowed to feed *ad libitum* on the inoculum for 2.5 hours at 28-30°C. An inoculum containing viable bacteria typically contains 10^3^-10^4^ colony forming units per µL (CFUs/µL), which was confirmed by dilution plating the inoculum on NA just before and just after the inoculation period. After 2.5 hours, nymphs were transferred to another sterile rearing box and starved for an additional 20-24 hours without water. This second starvation period was necessary not only to synchronize bouts of feeding activity among all nymphs in each experimental cohort but also to eliminate any nymphs that may not have ingested the inoculum.

For the second round of inoculation, liquid cultures were grown and washed as described above, using the same bacterial colonies that were used for the first round of inoculation. Inocula were prepared by diluting 5 µL of washed cells into 495 µL of inoculation medium, then dilution plated to confirm cell density, as above. To maximize exposure to the second strain, nymphs were exposed to this second inoculum for 24 hours before being individually isolated in 24-well cell culture plates with pieces of organic zucchini.

Nymphs were allowed to feed and develop on zucchini for 4-6 days after the second round of inoculation before sacrifice. Nymphs were killed with 70% denatured ethanol, and intact nymphs were imaged on an Olympus SZX16 stereomicroscope with an Olympus XM10 monochrome camera and Olympus cellSens Standard software ver. 1.13, as described in (35). Nymphs were immersed in a shallow volume of PBS in 6 cm plastic Petri dishes, and images were taken in darkfield (30 ms exposure 11.4 dB gain), brightfield (autoexposure, 11.4 dB gain), a GFP channel (autoexposure, 18 dB gain), and an RFP channel (autoexposure, 11.4 dB gain). Darkfield and brightfield images were merged in FIJI version 1.54f using the Image Calculator plugin, and the result was then merged with the GFP channel, RFP channel, or both.

To confirm imaging data, relative strain abundance within each individual was measured. After imaging, cadavers were surface sterilized in 10% bleach for 5-10 minutes, washed off again in 70% ethanol, and immersed in ∼20 µL droplets of PBS. M4s were individually dissected and stored in 300 µL PBS. Once all M4s from a set of nymphs were collected, samples were crushed with micropestles and dilution plated on NA. Plates were incubated at 30°C for 22-24 hours, then stored at 4°C until GFP and RFP colonies were enumerated.

### C) Antibiotic clearance experiments

The protocol for curing nymphs of symbionts is depicted in Figure 2A. Second instar nymphs were either inoculated with trimethoprim (Tp)-sensitive GA-OX1 RFP or with sterile inoculation medium for 2.5 hours, following the protocol as described above. The next day, nymphs were moved to a clean rearing box and given 200 µL of sterile inoculation medium containing either 167 µg/mL Tp, a canonically static antibiotic against which *Caballeronia* strains are extremely sensitive, or containing 200 µL of sterile inoculation medium amended with 20 µL of DMSO (solvent-only control). This procedure was repeated with a lower dose of 111 µg/mL Tp every ∼24 hours for the next two days due to antibiotic toxicity. After three days of Tp administration, all nymphs were allowed to feed on surface-sterilized organic zucchini for five hours to replenish their food reserves before being fed sterile inoculation medium containing 111 µg/mL Tp or DMSO for one more day.

After four days of Tp administration, nymphs were transferred to parafilmed, surface-sterilized organic zucchini pieces for the next three days to assess whether GA-OX1 RFP could resume proliferation within the M4 symbiotic organ during this time interval. At the end of the third day of recovery (one week after GA-OX1 was first administered), bugs were anesthetized with dry ice, immersed in sterile DI H_2_O, and their ventral aspects imaged as described above to confirm that GA-OX1 RFP was successfully depleted.

To qualitatively demonstrate the degree to which GA-OX1 RFP densities were reduced within Tp-fed insects, we also dissected the symbiotic organs from a subset of nymphs and examined them under a widefield scope. Dissected M4 organs were fixed with 4% PFA PBS, washed with PBS three times, permeabilized using 0.1% Triton-X PBS, washed with PBS three times, stained using 1:200 Phalloidin and 1:1000 DAPI overnight at 4 degrees Celsius, washed three times in PBS, and mounted using MWL-4 and AF300. Widefield images were acquired under 20X magnification with 200ms exposure time.

### D) Symbiont replacement experiments

To generate a trimethoprim-resistant, RFP symbiont (dsRed-TpR), electrocompetent GA-OX1 cells were first prepared by inoculating 1 mL of LB with a single colony and growing at 25°C overnight shaking at 200 rpm. The log phase culture was transferred in its entirety into 100 mL of yeast glucose medium (5 g/L yeast extract, 4 g/L glucose, 1 g/L sodium chloride) and grown at 30°C for 6 hours shaking at 225 rpm. Cells were gently pelleted for 5 minutes at 6,000 x g and washed in ice-cold 10% glycerol four times before being frozen at −80°C as 80-120 µL aliquots. The broad host range, multicopy plasmid pIN63 (45), which carries a Tp-resistant dihydrofolate reductase and a dsRed.T3 fluorescent protein, was introduced into GA-OX1 by electroporation of electrocompetent 40 µL of cells with 100 ng of plasmid DNA in a 0.2 cm cuvette on a Bio-Rad Micropulser (1 pulse at 3.0 kV voltage). Cells were recovered in 1 mL of yeast glucose broth and incubated at 30°C for two hours before plating on NA containing 0.4 µg/mL Tp for selection.

The protocol for Tp-sensitive symbiont replacement with the Tp-resistant strain is depicted in Figure 3A. Second instar nymphs were either inoculated with the Tp-sensitive GA-OX1 GFP or with sterile inoculation medium for 2.5 hours, then either fed with a Tp-laced or a control inoculation medium before recovering on organic zucchini as previously described. At the end of the third day of recovery, nymphs were starved for 12 hours before being fed GA-OX1 dsRed-TpR. Nymphs were then allowed to develop for a further week before imaging (as described above) to confirm GA-OX1 TpR colonization.

### E) Quantification of symbiont-elicited bottleneck remodeling in the constricted region

Nymphs were dissected and the gut fixed as described above. In images of the constricted region, measurements of the inner diameter of the constricted region were taken at the most anterior stretch using Phalloidin staining of microvilli as the marker for the epithelial cell apex. If the constricted region had completely closed off in at least one point, then the diameter was counted as 0.

### F) Statistical analyses

Differences in the relative abundance of RFP CFUs recovered from homogenized M4s were evaluated with a two-sided Wilcoxon test. The effect of Tp on GA-OX1 dsRed-TpR colonization of nymphs that had previously been inoculated with GA-OX1 GFP was assessed with a binomial regression, with Tp administration and GA-OX1 GFP presence as factors. To assess the change in the minimum diameter of the lumen within the constricted region, we used a one-sided Wilcoxon test.

## Acknowledgements

We thank Cristian Crisan (Goldberg lab) for providing the plasmid pIN63 and Sandra Mendiola and Erik Edwards for their assistance with the maintenance of insect colonies. We also thank Kota Ishigami (Kikuchi lab) for providing an electroporation protocol that we adapted for use in our experiments. Funding provided by USDA NIFA 2019-67013-29371.

---

**Figure S1.**
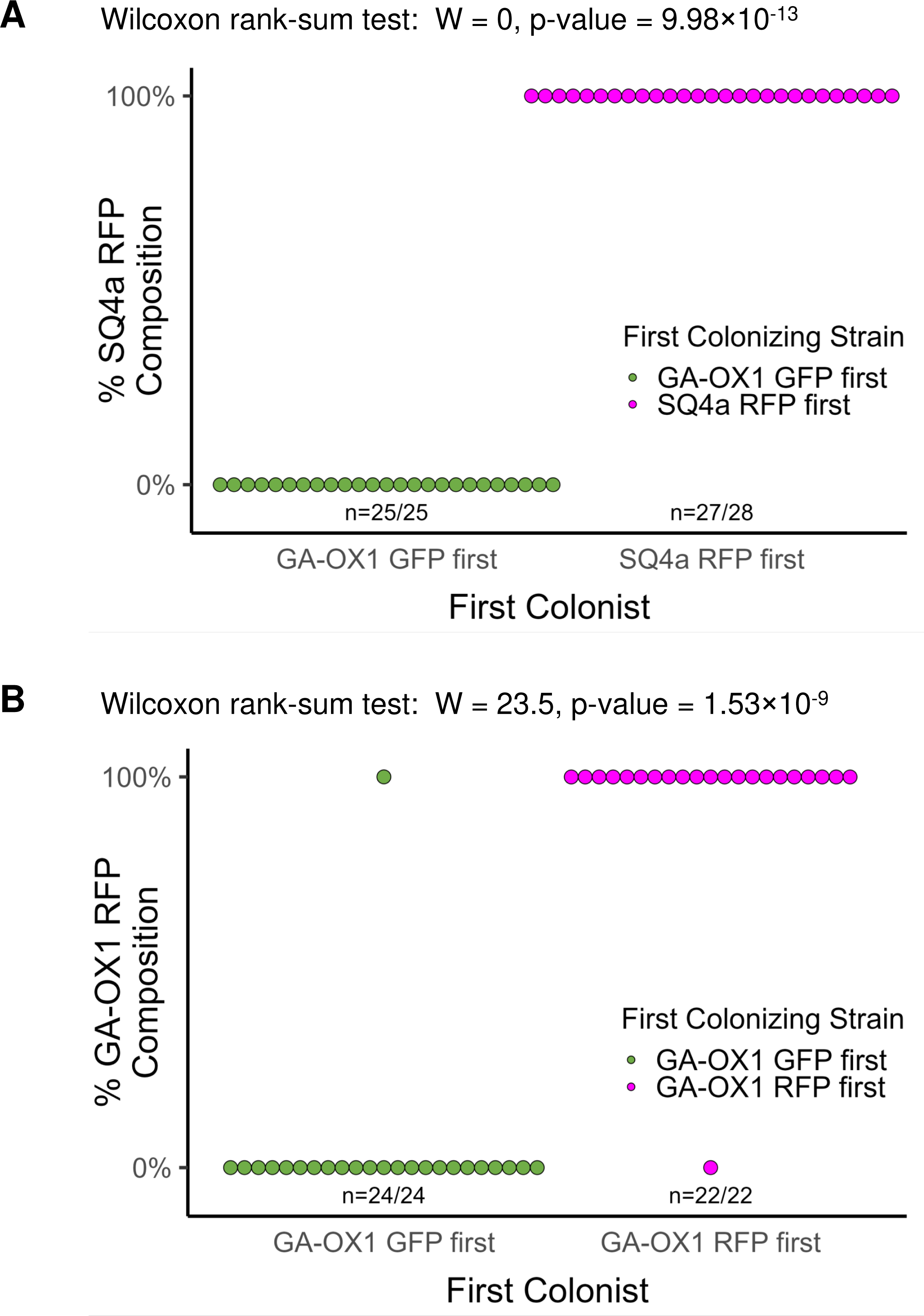
Relative abundances of unrelated and isogenic symbiont strains after sequential colonization. A) Relative abundance of GA-OX1 sfGFP and SQ4a RFP in squash bug nymphs after sequential colonization. Nymphs from which titers of each strain were obtained are depicted in Figure 1B. B) Relative abundance of GA-OX1 sfGFP and GA-OX1 RFP in squash bug nymphs after sequential colonization. Nymphs from which titers of each strain were obtained are depicted in Figure 1D.

